# Dynamic positioning and precision of bistable gene expression boundaries through diffusion and morphogen decay

**DOI:** 10.1101/2020.12.17.423202

**Authors:** Melinda Liu Perkins

## Abstract

1

Traditional models for how morphogen gradients guide embryonic patterning fail to account for experimental observations of temporal refinement in gene expression domains. Dynamic positional information has recently emerged as a framework to address this shortcoming. Here, we explore two central aspects of dynamic positional information—the precision and placement of gene expression boundaries—in bistable genetic networks driven by morphogen gradients. First, we hypothesize that temporal morphogen decay may increase the precision of a boundary by compensating for variation in initial conditions that would otherwise lead neighboring cells with identical inputs to diverge to separate steady states. Second, we explore how diffusion of gene products may play a key role in placing gene expression boundaries. Using an existing model for Hb patterning in embryonic fruit flies, we show that diffusion permits boundaries to act as near-traveling wavefronts with local propagation speed determined by morphogen concentration. We then harness our understanding of near-traveling fronts to propose a method for achieving accurate steady-state boundary placement independent of initial conditions. Our work posits functional roles for temporally varying inputs and cell-to-cell coupling in the regulation and interpretation of dynamic positional information, illustrating that mathematical theory should serve to clarify not just our quantitative, but also our intuitive understanding of patterning processes.

**Author Summary:** In many developmental systems, cells interpret spatial gradients of chemical morphogens to produce gene expression boundaries in exact positions. The simplest mathematical models for “positional information” rely on threshold detection, but such models are not robust to variations in the morphogen gradient or initial protein concentrations. Furthermore, these models fail to account for experimental results showing dynamic shifts in boundary placement and increased boundary precision over time. Here, we propose two theoretical mechanisms for enhancing boundary precision and placement using a bistable toggle switch. Distinct from existing research in “dynamic positional information”, this work posits a functional role for temporal decay in morphogen concentration and for diffusion of gene expression products, the latter of which is often omitted in quantitative models. We suggest that future research into dynamic positional information would benefit from perspectives that link local (cellular) and global (patterning) behaviors, as well as from mathematical theory that builds our intuitive understanding alongside more data-driven approaches.

## 3 Introduction

Spatial patterning of gene expression, especially during embryonic development, has fascinated theoretical and experimental researchers throughout the past century. A central question involves how an initial pattern is interpreted by cells in a tissue to produce sharp boundaries between spatial domains expressing distinct sets of genes [1], [2]. Significant evidence indicates that concentration gradients of chemical morphogens provide the necessary “positional information” for accurate boundary formation in various contexts ranging from axis orientation [3]–[6] to neurogenesis [7], [8]. In early studies, cells were proposed to interpret positional information through a “thresholding” mechanism, whereby a cell expresses a certain gene only if the morphogen concentration in that cell exceeds a certain level [9]. Subsequent research has complicated this hypothesis, with experimental evidence indicating that features such as the slope or temporal evolution of a morphogen gradient may also transfer positional information to target cells [10]–[16].

The recent concept of “dynamic positional information” posits that positional information is not conferred through a static “reading out” of morphogen concentration, but rather through a dynamic process of interplay between morphogen concentration and the genetic networks that interpret it [17]. Dynamic positional information has been inspired in part by two observations. First, the precision (regularity) of boundaries increases over time, which in combination with domain shifting leads to evolution of patterns from an initial “messy” form to a refined “final” form that cannot be directly predicted from the concentrations of underlying signals [18]–[20]. Second, gene expression domains shift over time, as indicated by experimental evidence from fruit flies [18], [21]–[25], scuttle flies [26], and vertebrate neural tube [27]. To explain these observed behaviors, researchers have investigated avenues including stochastic influences [28], [29], the structure of biochemical networks [30], protein diffusion [20], and the temporal evolution of upstream patterns [15].

Dynamical systems analysis—in which differential equations describe how chemical concentrations evolve in time—underlies most efforts to understand dynamic positional information. In a dynamical systems model, morphogen concentration typically acts as a spatially varying bifurcation parameter that modulates the number and stability of the steady states attainable by a genetic network (and therefore by a cell located at a particular point in space). The morphogen modulates gene expression rates such that cells destined for one side of a gene expression boundary favor one stable state and cells on the other side favor another stable state. Such models in the literature often neglect cell-to-cell (nucleus-to-nucleus) coupling through, for example, the diffusion of gene products, which simplifies simulation and analysis without completely compromising qualitative theoretical-to-empirical fit [15], [21], [30]–[32]. However, diffusion is known to be a potential determinant of patterning in chemical and biological systems alike—as demonstrated by Turing’s famous observation [33]—and theoretical and experimental research supports the idea that such intercellular signaling may reduce noisy influences or correct patterning defects in developmental systems [20], [34], [35].

Here, we explore two of the key elements behind dynamic positional information, precision enhancement and boundary shifting, using a simple model system, the bistable genetic network (“toggle switch”), which has previously been employed to model gene expression over morphogen gradients (e.g., [29], [35], [36]). First, we introduce the basic dynamical systems tools and terminology that we will use throughout the paper, in the context of a proposed mechanism for enhancing boundary precision. Specifically, we hypothesize a functional role for temporal decay of a morphogen gradient. The decay causes a bistable region flanked by two monostable regions to shift spatially in time, such that cells in initially bistable regions—where the boundary would be least precise in a system with a static gradient—are ultimately “swept up” by monostable regions. At the same time, “preinduction” toward a monostable state prevents cells that later become bistable from generating new irregularities in the boundary, resulting in a sharp and precise transition across the whole embryo. This “bistable sweep” also induces an apparent shift in boundary location as a consequence of the spatial shift in the bifurcation points.

Second, we analyze the effect of diffusion on the dynamic placement of gene expression boundaries. We consider a boundary as a near-traveling bistable wavefront, with the local front speed determined by the concentration of morphogen(s). We introduce the key concepts by examining an existing model for Hb boundary formation in early *Drosophila* [37], which exhibits both spatial and temporal variation in the underlying Bcd morphogen gradient. We then demonstrate the utility of the approach by analyzing a hypothetical robust mechanism for accurate steady-state boundary placement even from homogeneous initial conditions. In both this and the previous example, our analysis subtly shifts focus from individual cell trajectories to the evolution of entire gene expression boundaries as influenced by the spatiotemporal behavior of bifurcations across a model tissue. Our work highlights possible roles for temporally varying inputs and cell-to-cell coupling in the regulation and interpretation of dynamic positional information, which we believe to operate on both local and global scales.

## 4 Results

### 4.1 Enhanced boundary precision through temporal gradient decay

The simplest models for dynamic patterning assume static morphogen gradients. Experimentally, however, these gradients have been observed to change in time [18], [38]–[40]. In particular, temporal decay in gradient amplitude has been suggested to play an essential role in properly positioning gene expression boundaries [15]. Here, we hypothesize that gradient decay could not only place boundaries, but also enhance boundary precision by increasing robustness to variation in initial conditions that might otherwise cause neighboring trajectories to diverge to different stable steady states.

In this and the following section, we will approximate a tissue as a continuum, with dynamical behaviors varying along a coordinate *x*. For the sake of discussion we will take *x* to be the anterior-posterior axis of a model embryo. Assuming for now that parameters are static (not time-varying), we consider chemical reactions (without diffusion) governed by equations of the form

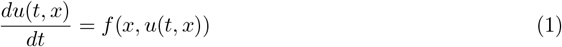

where *u*(*t, x*) are chemical concentrations at time *t* and coordinate *x*, and *f* incorporates the production and degradation rates of the chemicals. The whole system is presumed to be mono- or bistable at all points on the axis. For example, in this section we will consider a chemical “toggle switch” in which there are two mutually repressive species *u*_1_ and *u*_2_. The species *u*_1_ is activated by morphogen *α*, which is expressed in a gradient along *x* with length constant λ:

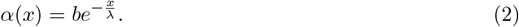

For simplicity, we also lump transcription and translation into a single step. The governing equations are then

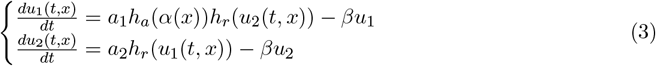

where

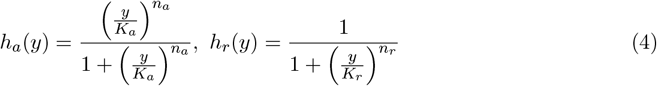

are activating and repressing Hill functions with coefficients *n_a_, n_r_* and concentration to half maximum *K_a_, K_r_* respectively.

The simplest boundary-forming system transitions from monostable to bistable back to monostable along *x* due to the spatial decay in the bifurcation parameter *α*(*x*). Sufficient variation in initial conditions will cause neighboring cells in the bistable region to diverge to separate stable steady states, such that in a two-dimensional embryo the resulting gene expression boundary will be imprecise (Fig. 1a).

**Figure 1:**
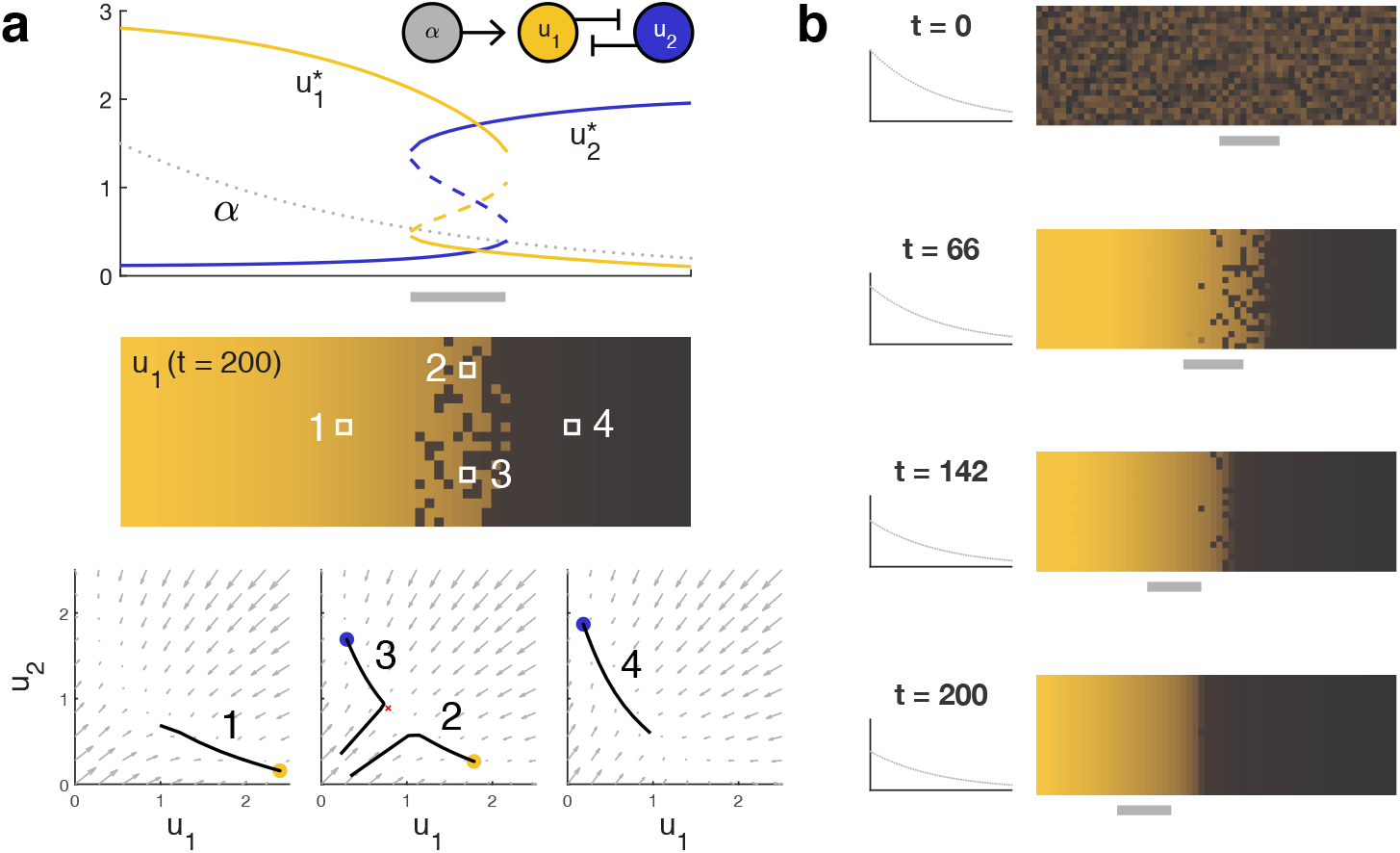
**(a) A bistable toggle switch can produce a gene expression boundary from a morphogen gradient.** Top, the morphogen (gray) induces expression of *u*_1_ (gold), which is mutually repressive with species *u*_2_ (blue). The morphogen concentration causes the anterior (left) of the boundary to be monostable in favor of high *u*_1_ and the posterior (right) to be monostable in favor of high *u*_2_. In the bistable region of the boundary itself, the initial conditions determine whether cells express high *u*_1_ or high *u*_2_ at steady state. Top, a bifurcation diagram indicates the value of the steady states, with solid lines indicating stability and dashed lines, instability. Middle, a heat map of *u*_1_ concentration at steady state showcases how variation in initial conditions among rows may result in an imprecise boundary in the bistable region. Bottom, phase portraits at different *x* coordinates along the embryo predict how trajectories evolve over time. Gray arrows indicate the direction in which a trajectory will travel. Solid dots indicate stable steady states favoring high *u*_1_ (gold) or high *u*_2_ (blue). The unstable middle point in the bistable region is denoted by a red *x*. Black lines are example trajectories corresponding to the cells labeled above. **(b) Temporal decay in the morphogen concentration at every point in space can “sweep away” imprecision in the expression boundary.** Decay causes the bistable region (solid gray line) to shift anteriorly over time. Thus, cells to the anterior are initially monostable for high *u*_1_ and remain there as they become bistable, whereas cells in the initially bistable region become monostable for high *u*_2_. The decay must be sufficiently slow for cells at the anterior to preinduce to high *u*_1_; see text and Appendix 8.

We will now consider morphogen gradients that are not static. Specifically, we will introduce time dependence to *α*(*x*) as

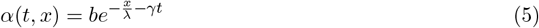

where *γ* is the temporal decay rate. More generally, we can rewrite the governing equation (1) as

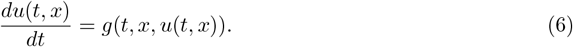

Cell behavior can thus be both local (fixed *x*) and instantaneous (fixed *t*). An instantaneous phase portrait corresponding to a fixed time *t* is derived from the equation

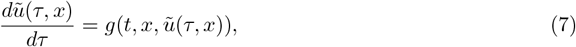

where the only difference from (6) is that *g* is fixed in time. For our purposes this corresponds to the time evolution of a hypothetical system in which the morphogen gradient is static, but with a profile corresponding to the profile of the time-varying system at time *τ*. For a more general discussion of instantaneous phase portraits, see [15].

In our toggle switch model with *α* decaying in time at all points in space, the transition areas between monostable and bistable regions shift anteriorly (toward low *x*) over time, such that parts of the initially monostable region for high *u*_1_ will become bistable, and part or whole of the initially bistable region will become monostable for low *u*_1_. If the decay is slow enough relative to the system dynamics, we can expect trajectories beginning in the anteriormost region to converge to high *u*_1_ while that region is monostable and to remain at the stable high *u*_1_ state as parts of that region become bistable with decaying *α*. At the same time, the initially bistable region becomes monostable for low *u*_1_, thereby “sweeping away” irregularities introduced by the initial bistability. The result is that an imprecise boundary can be made precise through the decay of the morphogen gradient (Fig. 1b). A side effect of this scheme is that the placement of the boundary, as determined by the locations of the bifurcation points, also shifts anteriorly. (Note that a similar effect could be observed but with a posterior shift if the morphogen concentration were increasing in time across the domain.)

In the classification scheme proposed by Verd et al. [41], trajectories on the posterior side of the boundary will be topologically captured by the low stable state as the high stable and middle unstable states disappear, regardless of their inital condition. On the anterior of the boundary, trajectories “pursue” the high *u*_1_ stable state even as topological changes introduce a low *u*_1_ stable state. Equivalently, the requirement for these cells is to evade topological capture by the low stable state as it emerges. Therefore, the “rate-limiting step” is the preinduction of anterior cells toward the high stable state; that is, the gradient should decay slowly enough in time that anterior trajectories are able to evolve “close enough” to the high stable state to remain in the basin of attraction as the bifurcation occurs to introduce the other two steady states. These effects are illustrated in Appendix 8 using a one-species, one-morphogen model with no time dependence.

### 4.2 Boundary shifts induced by near-traveling fronts

We now turn our attention to systems in which chemicals are permitted to diffuse. We will continue to use the terminology of local mono- and bistability established in the previous section, although in the presence of cell-to-cell coupling the trajectories of individual cells can no longer be predicted from local phase portraits alone.

It is well known that bistable systems with diffusion and spatially homogeneous parameters admit traveling wavefront solutions; biological examples include axonal propagation [42], population genetics [43], cell polarization [44], and Fgf8/RA gradient formation [45]. The speed and direction of front propagation depend on parameter values, specifically on which of the two stable steady states is more favorable. We propose that spatial variation in bifurcation parameters can be understood to modulate the local front velocity, providing a qualitative tool for analyzing how a front, once formed, will propagate along a domain. Local stability properties allow us to qualitatively predict where fronts emerge in response to arbitrary initial conditions. In this way, transient patterning behaviors can be analyzed apart from the trajectories of individual cells, with diffusion providing the link between local and global dynamics.

In this section, we will consider the counterpart to (1) with diffusion:

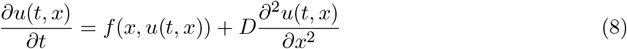

We will continue to refer to the local mono-or bistability of this system as the solutions at a particular coordinate *x* to the corresponding system without diffusion. At each coordinate we will also assign a local front velocity, which is the velocity of a traveling front solution to

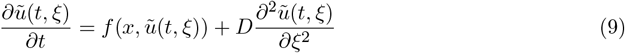

where *x* is fixed and *ξ* is a dummy spatial coordinate. In other words, the local front velocity captures the speed and direction that a traveling front would propagate if the parameters were spatially homogeneous at the values they assume at point *x* in (8). Analogously to the relationship between the time-independent dynamical system (1) and the time-dependent one (6), we can also add time dependence to (8) by replacing *f*(*x, u*) with some time-varying function *g*(*t, x, u*). In this case, the local front velocity is also instantaneous to a fixed time *τ* (corresponding to solutions to (9) with 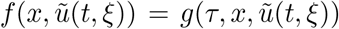). Throughout this section, we will use the term “near-traveling front” to distinguish shifting boundary solutions to (8) from true traveling fronts, which are translation invariant (i.e., their shape does not vary as they propagate).

#### 4.2.1 Example: Hb patterning over Bcd gradient in embryonic fruit flies

In a recent paper, Yang et al. quantified the anterior-to-posterior shift in the Hb boundary in wild-type, *stau*^−^ mutant, and half-Bcd dosage (Bcd1.0) mutant *Drosophila melanogaster* embryos [37]. Here, we analyze their model to reveal that the Hb patterning system is split between a locally monostable anterior and a locally bistable posterior with an extremely low front speed. Analysis through our framework clarifies the grounds for behavioral differences between *stau*^−^ and Bcd1.0 mutants, providing a more solid theoretical foundation for the authors’ observations.

Hb patterning is modeled as a one-dimensional reaction-diffusion system with no-flux boundary conditions:

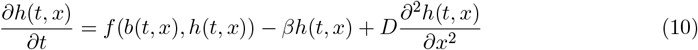

where *h*(*t, x*) and *b*(*t, x*) are the concentrations of Hb and Bcd respectively at A-P axis coordinate *x* and time *t, β* is the decay rate of Hb, and *D* is the diffusion constant of Hb. The regulatory function

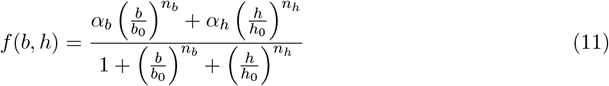

describes the activation of Hb expression by Bcd and Hb.

Consider the following dynamical system corresponding to (10) without diffusion:

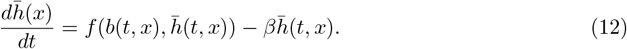

Further suppose that *b*(*t, x*) = *b*(*x*) is fixed in time. Then the Bcd concentration *b*(*x*) acts as a bifurcation parameter determining the local mono- or bistability of the corresponding reaction-diffusion system (10). Specifically, with all other parameters as given in [37], then with *b*(*x*) = 0 the system (12) is bistable, while for sufficiently high values of *b*(*x*) the system is instead monostable. We will denote by *b_crit_* = 0.07 the “critical concentration” where the system (12) transitions from bistable to monostable. An earlier model of Hb patterning also implicated Bcd in modulating bistability across the length of the embryo [36].

The Bcd gradient (with no temporal decay) is described by

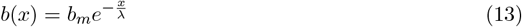

where *b_m_* controls the amplitude of the gradient and the length constant λ determines the dropoff. As long as *b_m_* > *b_crit_* (which is true in the authors’ three models), then the embryo will be locally monostable toward the anterior of the ambryo and locally bistable toward the posterior, with the transition occuring at the coordinate *x_crit_* where 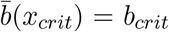. The bifurcation diagram reveals that the values of the high and low stable states in the bistable region are relatively constant some short distance from *x_crit_* owing to the low absolute change in *b*(*x*) as the gradient decays in space (Fig. 2a).

**Figure 2:**
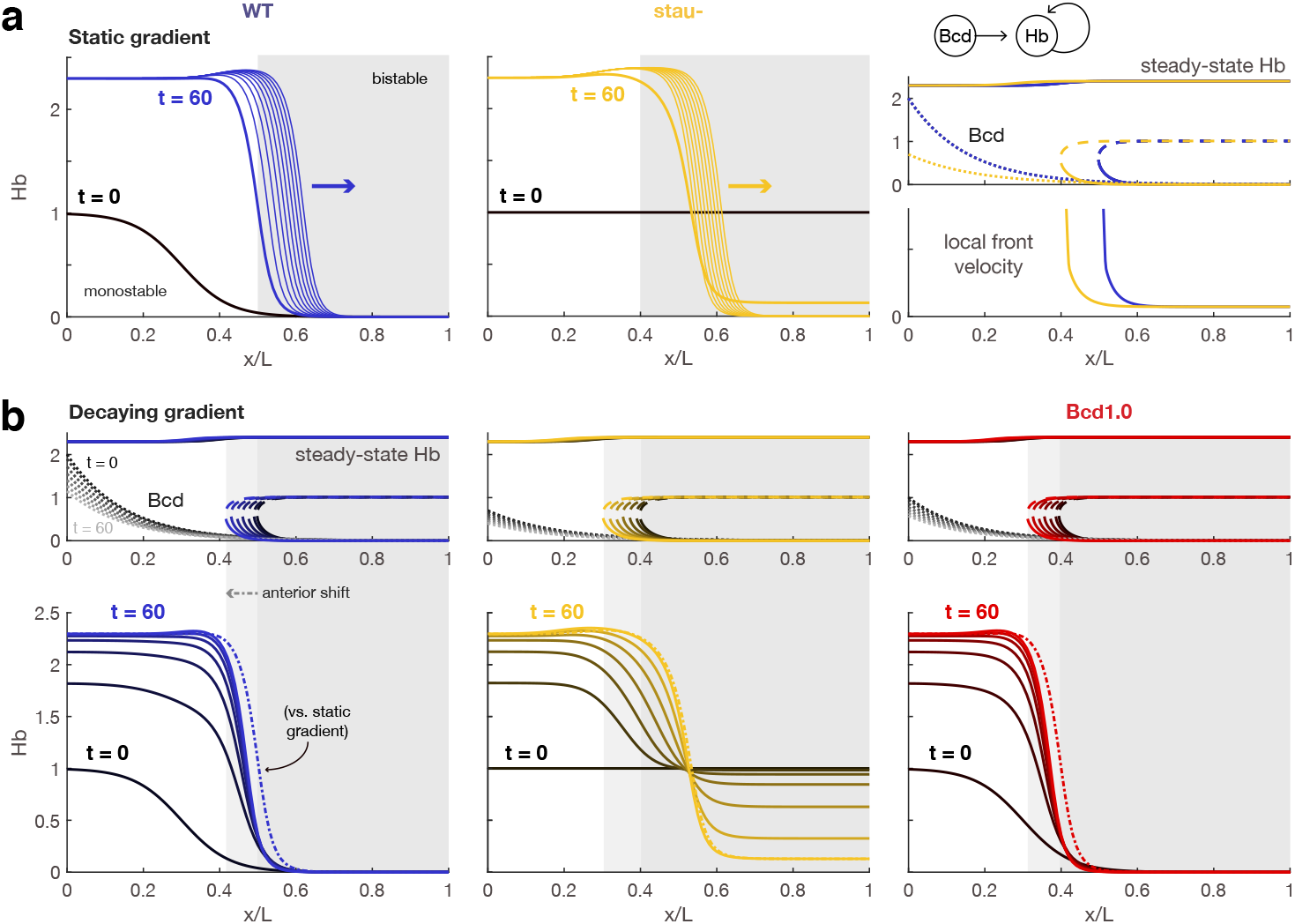
**(a) In the posterior of the embryo, the Hb boundary behaves as a bistable near-traveling wavefront.** Simulations (left, middle) and bifurcation diagrams (right) of the Hb patterning system with a static Bcd gradient show that the Hb gene expression boundary, which is “stabilized” around 60 minutes after nc14, in fact becomes a near-traveling bistable wavefront toward the posterior of the embryo (gray shaded region) and would continue to propagate slowly thereafter (thin lines) if not for external influences. Blue indicates wild-type (WT) and gold, *stau*^−^ mutant. Right, the Bcd gradient for *stau*^−^ (gold dotted line) has lower amplitude than that of the WT Bcd gradient (blue dotted line), which shifts the bifurcation (critical) point anteriorly. The local front speed rapidly decreases posterior to the critical point but remains positive, indicating that a bistable gene expression boundary will propagate posteriorly. The Bcd1.0 bifurcation diagram (see (b)) is almost identical to that of *stau*^−^. **(b) Decay in the Bcd gradient shifts the bistable region anteriorly over time, causing boundaries to stabilize at slightly anterior positions.** Bottom, there is a greater discrepancy between boundaries forming over decaying (solid lines) vs. static (dash-dot line) in the WT (blue) and Bcd1.0 (red) mutant than the *stau*^−^ (gold) mutant due to differences in front emergence. Specifically, homogeneous initial conditions below the maximum value of the unstable steady state cause the boundary in the *stau*^−^ mutant to appear within the bistable region, such that boundary propagation continues at roughly the same speed regardless of gradient decay. In contrast, in WT and Bcd1.0 the boundary is preinduced anterior to the critical point. Gradient decay causes the boundary to reach the critical point sooner, but also reduces the total time spent by the boundary in the region of rapid propagation, resulting in more anterior placement relative to the static case. All parameters are as given in [37].

The bifurcation diagram also shows that the switch is biased in favor of high Hb across the whole embryo. We therefore hypothesized that the Hb boundary, which was apparently “stabilized” 60 min after the start of nc14, was not, in fact, stable in the posterior of the embryo. To test this hypothesis we simulated (10) with *b*(*x*) fixed in space at values between 0 and b_crit_ corresponding to the values of *b*(*x*) with *x* > *x_crit_* in the model with varying Bcd gradient. From these simulations, we calculated the local front velocity *c*(*x*) at each coordinate *x* > *x_crit_*. We found that throughout the bistable region the local front speed is slow but nonzero (Fig. 2a), such that given enough time the near-traveling front would eventually vanish and leave high Hb in the entire embryo. This observation suggests that even with a static Bcd gradient the Hb pattern is in fact a long-lived transient solution, similar to those proposed to exist in a finite, spatially homogeneous domain [46]. This observation also accounts for the continued posterior migration of the boundary even after the transition width and steepness are fixed (Fig. 2a).

The local stability and front velocity are characteristics of the dynamical system determined by the choice of parameters. What behavior is actually observed, however, is also dictated by the choice of initial conditions. For example, the Bcd gradient is similar between *stau*^−^ and Bcd1.0 mutants, implying that the local dynamic properties are also similar. Thus, the authors differentiated between the two models by introducing distortion to the Hb initial condition ([37] Fig. 6D).

We now argue that we can qualitatively predict some system behaviors for arbitrary choices of initial conditions based on knowledge of the local dynamical properties. Specifically, in locally monostable regions of the embryo we expect the concentration to initially approach the local high state regardless of initial condition, while in locally bistable regions, we expect the concentration to initially approach the high state if the initial condition is locally above the unstable steady state, and to approach the low state if the initial condition is locally below the unstable steady state. Thus, in order for a Hb boundary to form over a Bcd gradient in this model, the initial condition must be such that the Hb concentration falls within the basin of attraction of the low stable state toward the posterior of the embryo, and within that of the high stable state toward the anterior of the embryo.

In WT and Bcd1.0, the initial condition is itself a boundary (with reduced magnitude) in the locally monostable anterior of the embryo. This initial boundary decays toward the value of the low stable state within the bistable region, such that in the absence of diffusion, the steady-state Hb in this region would remain at 0. The front is initiated by the diffusion of Hb produced in the monostable anterior into the bistable posterior. Halving Bcd concentration to produce Bcd1.0 shifts *x_crit_* to the anterior, thereby hastening the approach of the front to the locally bistable region where propagation is slow.

In contrast, in *stau*^−^ mutants the initial condition is homogeneous in space and falls just within the basin of attraction of the low stable state in the locally bistable posterior. Thus the boundary forms by the simultaneous diffusive influx of Hb from the monostable anterior and the attraction of Hb to the low state in the bistable posterior. Since the boundary evolves simultaneously toward both the high anterior and low posterior states (instead of beginning already at the low state in the posterior), the front is initially less steep than boundaries for the other two models. The large shift in Hb boundary position reported by the authors of [37] likely arises as a consequence of the “horizontal compression” of the boundary from an initially large and not very steep width to the sharp bistable front (Fig. 2b).

The discussion thus far has assumed no temporal decay in the Bcd gradient. Within our framework of near-traveling fronts, however, it is possible to incorporate the qualitative effects on system behavior. The authors model temporal Bcd decay through the addition of a decaying exponential term to (13):

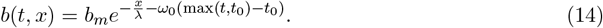

Such uniform-in-space decay of the Bcd gradient serves to shift *x_crit_* anteriorly over time. This hastens the “capture” of emerging fronts in the locally bistable region of slow propagation, thereby “stabilizing” the fronts at a slightly anterior position relative to predictions without gradient decay (Fig. 2b). Further research would be necessary to determine whether the temporal decay of the gradient plays a major role elsewhere in patterning, or arises purely as a consequence of other biological processes.

In summary, we found that the Bcd gradient acts as a spatially varying parameter that modulates the local steady-state properties of the reaction-diffusion system and, in locally bistable regimes, also modulates the speed and direction at which a bistable front will propagate. Initial conditions determine where the boundary emerges and therefore whether pattern “stabilization” is determined by the point at which the boundary reaches the locally bistable region (as for the WT and Bcd1.0 models) or by the convergence of the boundary to its final shape (as for the *stau*^−^ model). In addition to these insights, our perspective on expression boundaries as near-traveling fronts hints at classes of changes to the model that could better recapitulate experimental results. For example, any model components that tip the bistable system in favor of high Hb would increase the front speed in the posterior, thereby recapitulating the continued motion of the boundary observed in the Bcd1.0 model. In this case, possible factors could include additional network components or any modifications resulting in increased Bcd concentration in the posterior at later time stages (e.g., through flattening of the gradient over time).

#### 4.2.2 The final front(ier): Accurate boundary placement through front localization

In addition to offering insight into existing models, thinking about gene expression boundaries as near-traveling fronts reveals intriguing mathematical hypotheses for pattern-forming systems yet to be found in nature. Here, we consider the case where the local front velocity passes through zero at some critical point in the domain. This implies a switch in propagation direction, such that a front forming on either side of the point will propagate toward it. This scheme decouples the steadystate location of the boundary from the exact value of the homogeneous initial condition, and also produces qualitatively predictable transient behaviors. Once a localized boundary is established, its stability to intrinsic and extrinsic perturbation may be further analyzed to assess its reliability as a cue for downstream patterning processes [35], [47].

A number of developmental patterning systems employ opposing morphogen gradients, perhaps to enhance accuracy in boundary placement [48] or confer robustness to tissue scaling [9], [49]. We employ a simple model of a chemical “toggle switch” in which two mutually repressive species *u*_1_, *u*_2_ are activated by morphogens *α*_1_, *α*_2_ respectively. The morphogens are expressed in opposing gradients. For simplicity, transcription and translation are lumped into a single step. The governing equations are

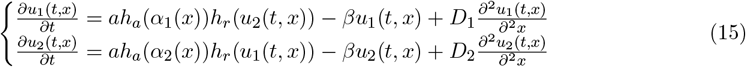

where *h_a_, h_r_* are activating and repressing Hill functions respectively, as in (4).

Assume *D*_1_ = *D*_2_ = *D*. The front localizes at the point *x_l_* where the switch is “balanced” such that neither of the two stable steady states is dominant, i.e., *α*_1_(*x_l_*) = *α*_2_(*x_l_*) (assuming *x_l_* is sufficiently far from the edges of the domain). Therefore, in this scheme, the final placement of the gene expression boundary relative to the domain edges will be unaffected by changes to *α*_1_(*x*) and *α*_2_(*x*) that do not shift *x_l_*, e.g., scaling the embryo length or equally modulating the amplitude or spatial decay rates of the gradients (provided that bistability is maintained in the region near *x_l_*). Quantitative features such as the boundary steepness and propagation speed may still depend on the exact parameter values.

From the bifurcation diagram, we can predict that uniform-in-space initial conditions will produce a front. Notice that here we have collapsed the effect of two bifurcation parameters, *α*_1_(*x*) and *α*_2_(*x*), onto a single bifurcation parameter, the axis *x*. The switch in the anterior of the domain is biased in favor of *u*_1_ while the posterior is biased in favor of *u*_2_, such that there is a range of concentrations for which initial conditions homogeneous or near-homogeneous in both chemicals will fall into the basin of attraction of *u*_1_ toward the anterior and *u*_2_ toward the posterior. This means that a properly oriented front (high *u*_1_ in the anterior) will emerge and approach the localization point. The initial location where the front appears depends on a combination of the numerical values of the initial concentrations, and the relative rates of front emergence (local divergence toward high or low steady state) vs. local front speed at the point where the initial concentrations cross from one basin of attraction to the other. In contrast, a “front” forming in a system without diffusion would remain at the initial location where it appeared, rendering the placement of the boundary more sensitive to initial conditions (Fig. 3a).

**Figure 3:**
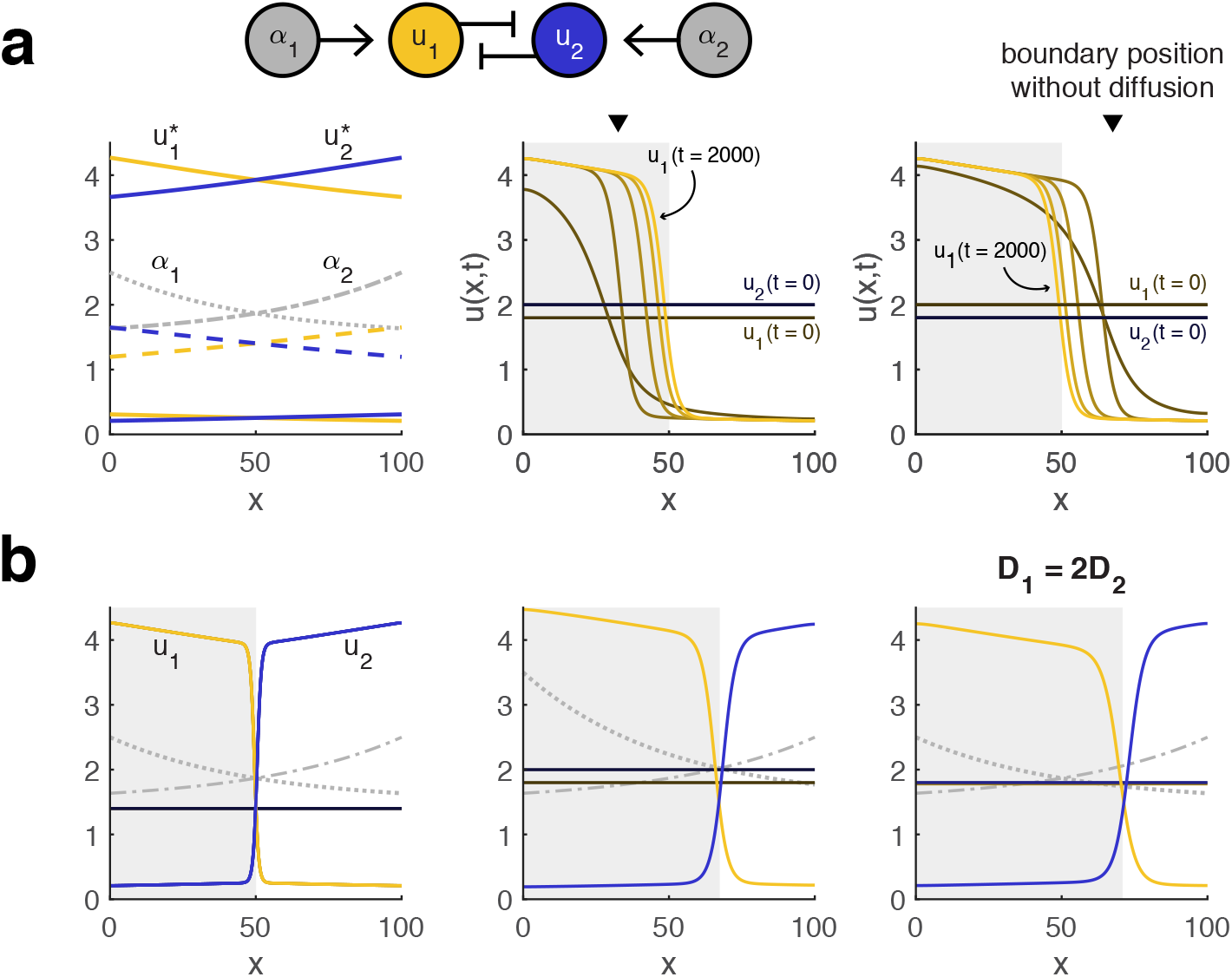
**(a) Diffusion permits a bistable gene expression boundary to be accurately placed independently of initial conditions.** Left, bifurcation diagram and right, simulation results for (15) with *a* = 1.7, *β* = 0.35, *D*_1_ = *D*_2_ = 1, *n_a_* = *n_r_* = 2, *K_a_* = 0.75, *K_r_* = 1, and *α*_1_(*x*), *α*_2_(*x*) as shown. Shaded region indicates rightward propagation of the front in *u*_1_ (gold) and leftward for the front in *u*_2_ (blue). Regardless of the initial condition, the boundary propagates toward the point where the toggle switch is “balanced” (the local steady states are such that 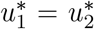). Inverted black triangle indicates the position of a gene expression boundary with the same initial conditions when gene products do not diffuse. **(b) Imbalance in gradients or diffusion may shift the location of the boundary.** Left and middle, when *D*_1_ = *D*_2_, the boundary localizes where *α*_1_(x) = *α*_2_(*x*). Right, letting *D*_1_ = 2 = 2*D*_2_ shifts the localization point in favor of higher *u*_1_. All other parameters are the same as for (a).

If *D*_1_ and *D*_2_ are unequal, then the localization point no longer corresponds to the point where the switches are locally balanced. Specifically, the localization point 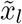 shifts to favor the faster-diffusing species, with larger disparities increasing the size of the shift. Thus, differences in diffusivity reverse the direction of front propagation in the region between *x_l_* and 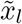 (Fig. 3b). Recent research into cell polarization indicates that two mutually repressive species with highly disparate diffusivities may also produce bistable fronts that are fixed in place. However, this *wave pinning* mechanism is distinct from front localization in that the highly diffusive species acts as the bifurcation parameter, and its decay over time “pins” a propagating boundary in place [44], [50], [51]. In other words, propagation is halted through the temporal modulation of a homogeneous-inspace parameter controlling the front velocity, rather than through spatial variation of exogenous parameters that locally bring the front velocity to zero.

Future quantitative investigation of developmental biological systems may reveal whether the front localization mechanism occurs in natural contexts. In the meanwhile, an understanding of near-traveling fronts can inform attempts to engineer synthetic multicellular patterning. For example, researchers have recently implemented patterning over concentration gradients with bistable networks in bacteria [52], [53]. Our results suggest that adding diffusion of gene expression products to such systems could enable dynamic, predictable boundary placement without the need to manipulate bifurcation parameters in real time. We may also envision more complicated patterning schemes for natural or synthetic application, particularly in systems where a single domain possesses multiple localization points (Fig. 4).

**Figure 4:**
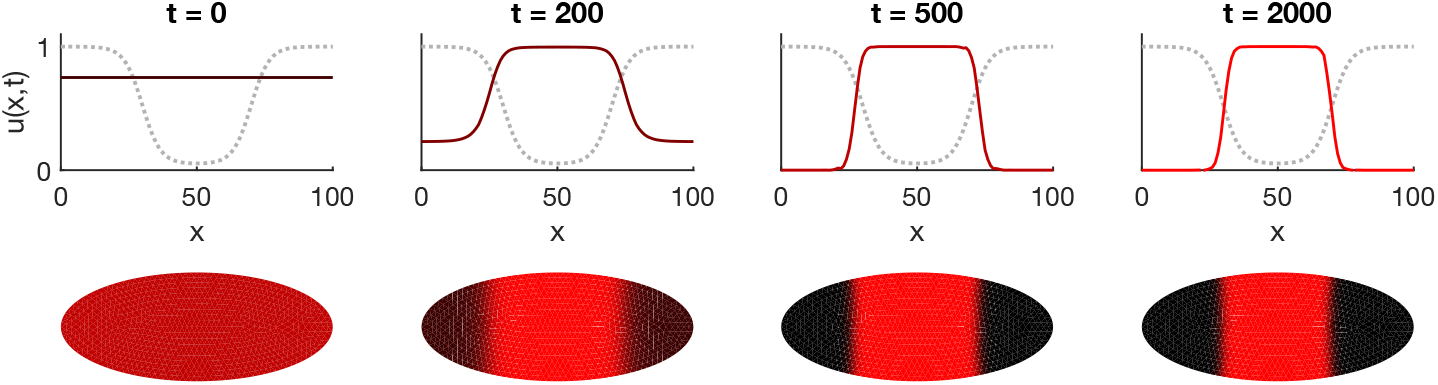
Fronts may localize at multiple points in a domain, resulting in more complicated patterns. Simulations were formed on the 1D model in (22) with *a* = 0.1, *D* = 0.1 on a 2D reaction domain (bottom) in which the bifurcation parameter *α*(*x*) varies only along the horizontal axis (gray dashed line). Top, horizontal cross-sections through the midline show that two opposing fronts appear and propagate toward the nearest localization point, resulting in refinement of a stripe pattern from either direction over time.

## 5 Discussion

Traditional models for morphogen gradient interpretation are susceptible to variation in initial chemical concentrations and the shape of morphogen gradients [54] as well as to intrinsic stochasticity in gene expression [29], [35], [55], [56]. Dynamic positional information has been proposed as a framework to address the resulting irregularities in gene expression boundaries [56] and to better recapitulate empirical observations [17]. In this work, we have proposed mathematical mechanisms for boundary precision and placement from a standpoint that links individual cell trajectories to the behavior of entire patterning features. We have hypothesized a functional role for time-varying morphogen concentrations and have shown that incorporating diffusion into modeling reveals an alternative mechanism to account for experimentally observed shifts in gene expression domains.

Although we have taken a bistable switch as our conceptual model, we believe the basic ideas explored herein may be generalized to a broader class of systems. Our hypothesized mechanism for enhancing boundary precision essentially relies upon the ability of cells to be preinduced to a desired state before temporal changes introduce further bifurcations. Such (evasion of) topological capture as a means to correct for variation in initial conditions could generalize to more complicated multistable networks. In such cases it may be appropriate to draw upon mathematical tools from the growing literature on nonautonomous dynamical systems; for example, the notions of pullback attractors and tipping points could be employed to determine the maximum rate at which bifurcation parameters can evolve in time without disrupting the ability of a trajectory to track a desired steady state [57], [58].

Similarly, near-traveling fronts could be found in bistable systems of higher than two dimensions with many more bifurcation parameters than considered here. With appropriate care, bistable subsystems might even be dissected from more complex networks, with the contributions of those networks serving as time-varying parameters or as factors that place bounds on dynamic behavior. Traveling wavefronts in non-bistable systems are also known to exist, for example, in monostable systems, although unlike bistable fronts there is a minimum speed at which they must propagate [59]. Continuing mathematical investigation will doubtless open up new avenues for exploring near-traveling fronts in gene expression patterning and other areas of embryonic development [60].

Even working from a simple bistable model, our analysis highlights a few ways in which dynamic positional information may be further explored to better our understanding of gene expression patterning, both philosophically and mathematically. Consider signaling duration, which has emerged as a central feature of morphogen-signaling systems [14]. From a dynamical systems perspective we might expect signal duration to matter even with a static gradient: Supposing a signal modulates local mono- or bistability, then if a cell begins from a low initial condition, it may take longer to approach a high expression state than a low one simply because it takes more time to produce more protein than to produce less. This observation may partially explain some experimental results, e.g., that anterior patterning in embryonic *Drosophila* is more highly disrupted by transient perturbations to areas with high than low Bcd concentration [13]. Signaling duration also plays a role in our two hypothetical mechanisms. In the bistable sweep model, cells to one end of the boundary require a minimum amount of time to “lock in” to the appropriate steady state before a bifurcation arises elsewhere in phase space. As we found in our examination of near-traveling fronts, signaling duration in one region of a tissue can also determine fates in other regions of the tissue simply because a gene expression boundary requires time to propagate. In the case of Hb patterning, decreasing signal in the anterior of the embryo decreased propagation speed, thereby shifting the entire boundary location anteriorly.

In addition to explicitly incorporating temporal evolution, dynamic positional information has implicitly challenged the notion of “threshold-dependent activation”, which we argue conflates the influence of two factors: the morphogen and the initial condition. In the simple boundary-forming model in Fig. 1a, the genetic network does indeed act as a threshold detector for very high or low morphogen concentrations, since cells in these regions are monostable and therefore their final fates are independent of initial conditions. However, there is a band of concentrations for which the system is bistable and cells must be in a “responsive” state induced through initial conditions or coupling with neighbors in order to approach the appropriate final expression level. Thus, for a fixed uniform initial condition, the apparent threshold at which cells are activated will shift, giving rise to the observed inaccuracy in boundary placement in such models.

Local threshold-dependent activation also cannot explain boundary placement in the bistable sweep or front localization models, even though both exhibit predictable behavior regardless of initial condition. In the bistable sweep model as illustrated in Fig. 1, the sharp boundary is ultimately found at the posterior transition from bistable to monostable, which shifts anteriorly over time. In the front localization model, the boundary position is fixed across the whole range of initial conditions that produce a boundary, such that a “threshold” for activation is determined by the single point where the front speed crosses zero. In both cases, individual cellular trajectories have less predictive power for the final observed pattern than do global dynamical properties.

The matter of thresholding and initial conditions raises a further philosophical point in relation to the notion of cellular “competence”, or the ability to “[express] all the necessary components for receiving a signal” [14]. Consider again a bistable toggle switch of two species, *A* and *B*, in which expression of *A* increases in response to increasing morphogen concentration (but never to the point of monostability). An upstream network component that establishes high *A* or low *B* as an initial condition could be considered necessary for competence, insomuch as the system lacking this component would appear not to respond to signal (*A* might always end up at the low steady state). But this situation is clearly distinct from the loss of a regulatory interaction between *A* and *B*, which would destroy the ability of the cell to respond to signal regardless of initial condition. Of course, it is rare that real developmental systems decouple nicely into components that only establish initial conditions vs. those that govern subsequent dynamic evolution. Nevertheless, the distinctions made clear through a precise mathematical representation suggest potential areas to refine our qualitative understanding and intuition.

Taken together, our observations support the notion that positional information is neither locally contained nor statically interpreted. Dynamic features may play functional roles in enhancing patterning robustness to initial conditions. At the same time, tissue-level patterning behaviors may not be immediately predictable from examining individual cell trajectories, especially in the presence of cell-to-cell coupling. Future research in gene expression patterning will benefit from further techniques for modeling temporal evolution at multicellular scales, particularly with respect to transient dynamic behaviors. Whether the hypothetical mechanisms explored here are employed in natural embryonic development remains to be seen. Nevertheless, we are confident that their presence or absence will prove insightful in our quest to disentangle explanation from observation, function from form.

## 6 Methods

All simulations were performed in Matlab 2017b.

## 7 Acknowledgments

The author would like to thank Murat Arcak for suggesting a numerical method to verify stability of 1D localized front solutions in finite domains. The author also thanks Justin Yim for providing code to approximate the separatrix of a 2D bistable system, and Regina Eckert for providing code to generate tile plots.

## 8

## Appendix 1: 2D autonomous model for precision enhancement through gradient decay

In the following discussion, we consider the 1D time-dependent (nonautonomous) model

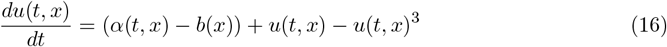

at coordinates *x* and times *t*. We assume there is some set of coordinates where the system is instantaneously bistable, with boundaries denoted by *x*_1_(*t*), *x*_2_(*t*); that is, *x* < *x*_1_(*t*) implies (16) is instantaneously monostable (high *U*) at time *t, x*_1_(*t*) ≤ *x* ≤ *x*_2_(*t*) implies (16) is instantaneously bistable at time *t*, and *x*_2_(*t*) < *x* implies (16) is instantaneously monostable (low *U*). Assume several realizations of the process are given, each ultimately representing a horizontal slice of a 2D “embryo”. For convenience we define 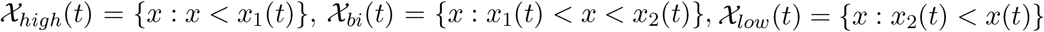, where the subscripts are derived with respect to the expected steadystate concentration of *U*.

We will consider the case where *α*(*t, x*) varies smoothly and monotonically in time, that is, *t*_2_ > *t*_1_ ⇒ *α*(*t*_2_, *x*) ≤ *α*(*t*_1_, *x*) ∀*x*, and asympototically approaches a finite value: *α*(*t* → ∞, *x*) → *α*\*(*x*). For most of this discussion we will assume *α*(*t, x*) decreases, although it is conceptually straightforward to generalize our discussion to the case where *α*(*t, x*) increases. For our setup, decay of *α*(*t, x*) in time means *t*_2_ > *t*_1_ ⇒ *x*_1_(*t*_2_) < *x*_1_(*t*_1_) and *x*_2_(*t*_2_) < *x*_2_(*t*_1_). We will further assume that 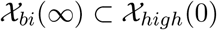 and 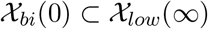, although this requirement may be relaxed.

If the decay of *α*(*t, x*) is much faster than the dynamics of *u*, then the behavior of the system is well approximated by (16) with *α*(*x*) = *α*\*(*x*). If the dynamics of *u*(*t, x*) are much faster than the decay of *α*(*t, x*) and the system satisfies certain technical requirements including slow evolution of *α*(*t, x*) [58], then we expect that *u*(*t, x*) will rapidly converge to the instantaneous system attractors and will in effect traverse the *stable paths* of those attractors. Thus, 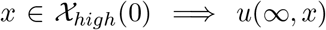 high, and 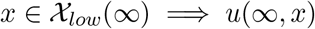 low regardless of initial condition. In other words, at infinite time there should be no remaining irregularity at the boundary between high and low *U*.

If the dynamics are on similar timescales, however, then variation in initial conditions may exert influence on system outcome: Two trajectories *u*(*t, x*) with 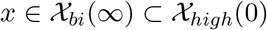 may converge to different steady states, such that the final boundary is imprecise. Guaranteeing a precise boundary, therefore, becomes a matter of restricting variation in initial conditions for 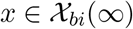 to the basin of attraction of the stable path corresponding to high *U*.

To examine this question intuitively, we now assume a particular form of *α*(*x, t*) that allows us to rewrite (16) as a time-independent (autonomous) system. In particular, suppose that the morphogen at coordinate *x* decays exponentially with rate *r*:

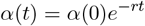

where for simplicity of notation we have dropped the dependence of *α* on *x*. The time evolution of *α*(*t*) may be described by

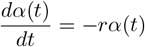

for initial condition *α*(0). We can then rewrite (16) as

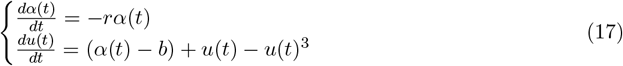

where 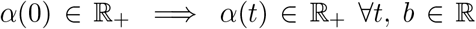, and we have again dropped the dependence of *u* and *b* on *x* for simplicity of notation. The system (17) has saddle-node bifurcations at each of 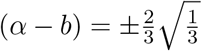 [61]. Since *α*(*t* → ∞) → 0, the steady states in this system are set by *b*:

- one stable high state if 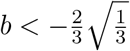,
- two stable states and one intermediate unstable state if 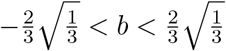,
- one stable low state if 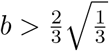.

In a toy model of an embryo, we would expect *r* to remain constant across the axis while *α*(0) and *b* form spatial gradients determining the initial morphogen concentration and mono- /bistability respectively. Since monostable systems have outcomes that do not depend on initial conditions, to ensure boundary regularity we must ensure that bistable cells end up in the high state. Specifically, we seek to determine for which range of initial morphogen concentrations *α*(0) and chemical concentrations *u*(0) trajectories terminate in the high state. This is the definition of the basin of attraction for the high state, represented in Fig. 5 by the region above the thick red line in each plot. Decreasing *b* or *r* increases the size of the basin of attraction for the high state, which matches the intuition that a longer time spent with a higher input ought to bias the system toward a higher state. As *r* increases, the system behaves effectively as though *α*(*t*) = 0, since trajectories evolve much more slowly than the change in the input level.

**Figure 5:**
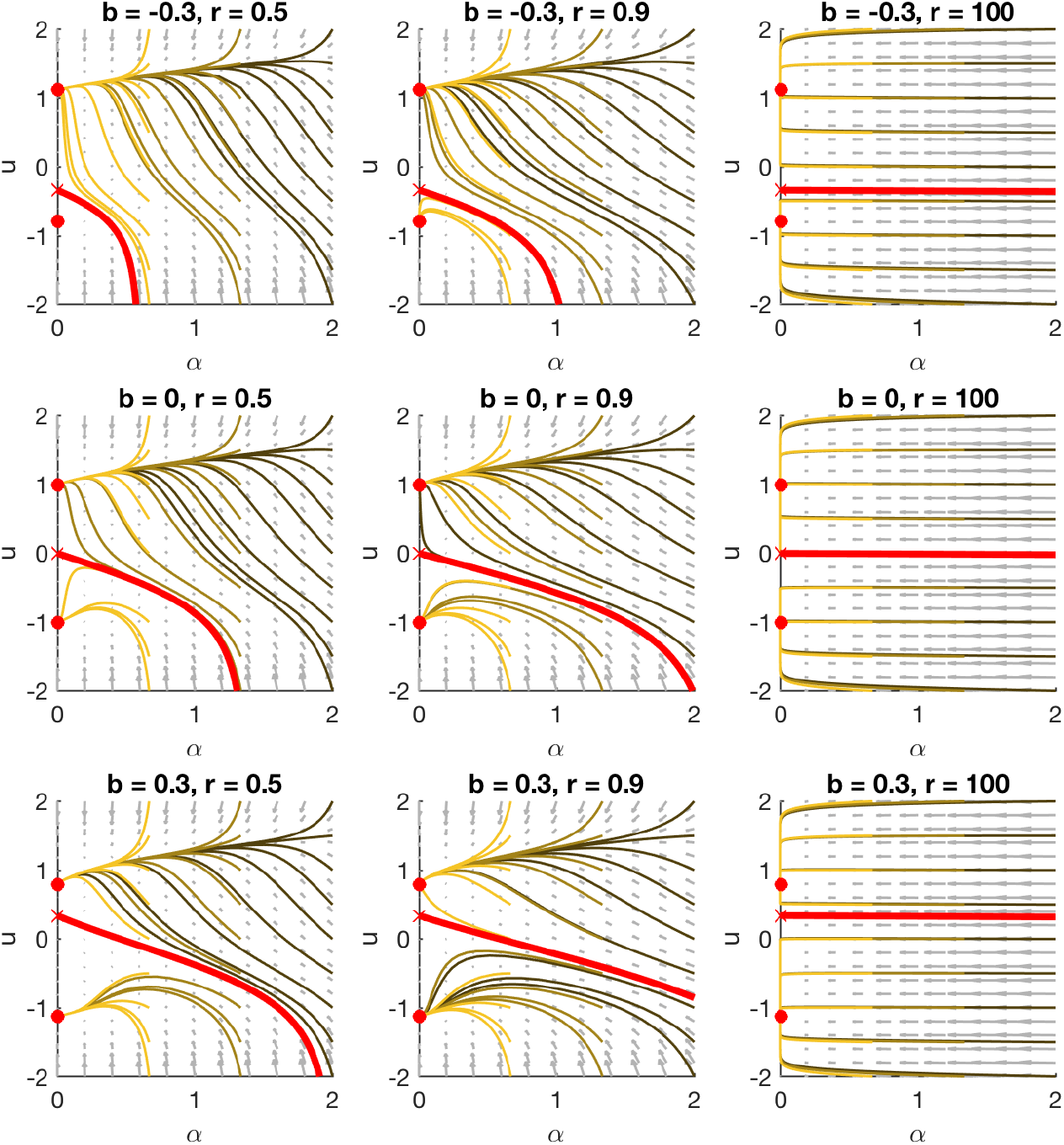
Decreasing decay rate causes more trajectories to reach the high steady state. Phase spaces (gray vector field) overlaid with sample trajectories (gold) for (17) with varying *b, r*. Darker trajectories start with higher *α*(0). Red circles denote stable stady states and the red *x* denotes the unstable steady state. The thick red line is the separatrix, so named because it separates initial conditions that end in the high state (above the line) from those that end in the low state (below the line). As *b* or *r* is decreased (to the top or left), the basin of attraction of the high state increases in size; i.e., trajectories starting with a wider range of initial conditions will end up in the high state. For fixed *b, r*, a boundary can form when the initial gradient *α*(0) spans the separatrix.

## Appendix 2: 1D model for front localization on a finite domain

Here we provide a more mathematical explanation for bistable front localization using a single-species model in a one-dimensional reaction domain. We consider systems of the form

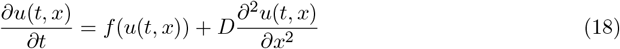

with 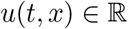 and

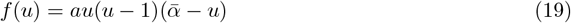

for 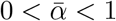. We assume *u*(*t, x*) ∈ [0, 1] ∀(*t, x*).

Suppose 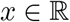, such that the reaction domain is an infinite line. Then the system (18) admits a traveling wavefront solution with profile 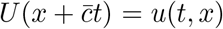. Analytical solutions are known for *f*(·) as in (19). In particular,

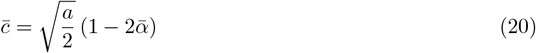

such that the wave ceases to propagate when 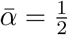 [42]. (Traveling fronts on infinite discrete lattices may localize for a wider range of parameters than corresponding continuous systems, depending, for example, on coupling strength [62] or propagation direction [63].)

Now consider the solution to (18) on a finite domain *x* ∈ [*x*_0_, *x*_1_] with no-flux (Neumann) boundary conditions

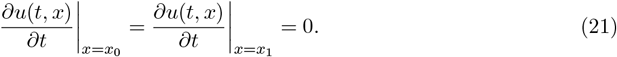

Although no stable, nonhomogeneous steady-state solutions exist for 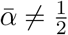 in the continuous case [64]-[66], transient (but potentially long-lived) wavefronts may emerge that propogate from one end of the domain to the other before vanishing [45], [46].

Researchers have previously noted that spatial variation in 2D domain shape [67] or reaction-diffusion terms (e.g., [68], [69]) can lead to stable nonhomogeneous solutions. We now introduce a class of spatial inhomogeneities that stabilize particular solutions comprising one or more localized fronts, where by “front” we mean an approximately sigmoidal curve. Let

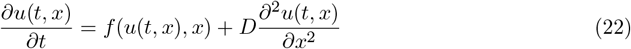

where

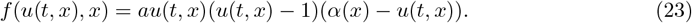

The dependence of *f* on *x* permits stable nonhomogeneous solutions to exist. We enforce that *α*(*x*) ∈ (0, 1) ∀*x*, such that *f* ∈ [0, *a*]. We further assume *α*(*x*) is smooth and monotonically decreasing, i.e., 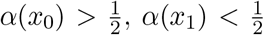, and 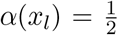 for some *x_l_* ∈ (*x*_0_, *x*_1_). We associate with each *x*\* ∈ [*x*_0_, *x*_1_] a local front velocity *c*(*x*\*), which is the velocity (20) of a monotonically increasing traveling wavefront solution to (18) for 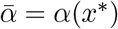.

Suppose *u*(*x*, 0) is a monotonically increasing front with transition width much smaller than the length of the domain. Since *α*(*x*) is defined such that *c*(*x*) < 0 for *x* < *x_l_* (rightward propagation) and *c*(*x*) > 0 for *x* > *x_l_* (leftward propagation), we would intuitively expect the front to approach the localization point *x_l_* where *c*(*x_l_*) = 0 regardless of where the front is located in the domain.

This observation suggests that there is a stable steady-state front solution centered at *x_l_*, which we numerically confirmed to be a stable nonhomogeneous solution in the discrete-space case. Note that these front solutions do not match the stable profile *U*(·) for any corresponding system with homogeneous 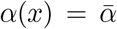. Reducing *D* reduces the boundary width, “sharpening” the transition between expression states (Fig. 6b).

**Figure 6:**
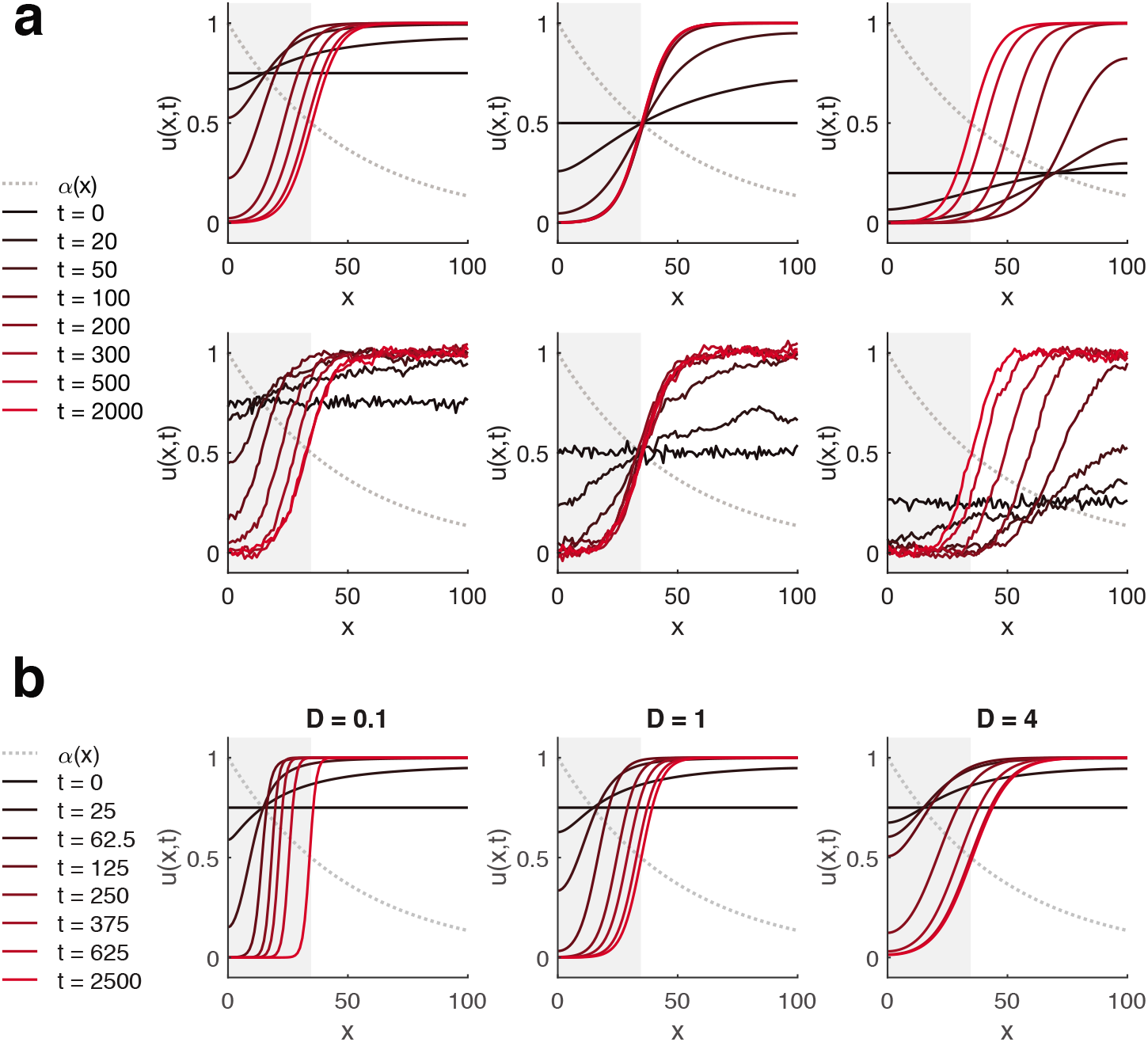
**(a) A near-traveling wavefront may localize to the same point in the domain regardless of uniform-in-space initial condition.** Localized front solutions to a 1D toy model (22) with *a* = 0.1, *D* = 2, and *α*(*x*) (gray dotted line) as shown. The value of a homogeneous initial condition (black line, *t* = 0) determines where the front emerges. Subsequent propagation (rightward in gray shaded region, leftward otherwise) causes the front to approach the localization point where the local front velocity is 0 (brighter red indicates later time points). Bottom, small amounts of noise do not disrupt the behavior, as suggested by stochastic simulation of a discretize-space model (*N* = 101 points) with Langevin noise of standard deviation *σ* = 0.02 (Euler-Mayurama method, step size Δ*t* = 0.2). **(b) Reducing** *D* **reduces boundary width.** All other parameters are the same as for (a). The local front velocity is independent of *D*, but in these simulations sharper boundaries appear to approach the localization point more slowly. We hypothesize this phenomenon relates to the dynamics of convergence to front profiles from homogeneous initial conditions.

A system with a localized front both admits a stable steady-state front solution and can reach that solution from spatially homogeneous initial conditions. Define for each *x* an equivalent system to (22) with no diffusion, governed by

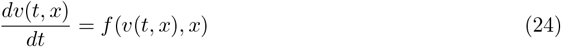

for *f*(·, ·) as in (23). At each point *x, v*(*x*, ·) has two stable steady states 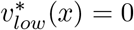 and 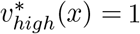. Let *v*(*t, x* = 0) = *b* ∈ (0, 1). For a given *x, b* will fall in the basin of attraction of 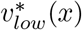 if *b* < *α*(*x*) and in the basin of attraction of 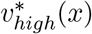 if *b* > *α*(*x*). In the corresponding diffusive system (22), the front will appear between *x_l_* and *x_b_* satisfying *b* = *α*(*x_b_*), with the location depending on the relative rates of front emergence (local divergence toward high or low steady state) and instantaneous front propagation. Thus, persistent spatial parameter variation produces a front with a particular orientation and ensures the convergence of that front to a fixed location in the domain, without requiring spatial bias in initial conditions (Fig. 6a).

